# Transcriptional repression by AP2-Sp3 regulates the mosquito-to-mammal infectivity switch in malaria sporozoites

**DOI:** 10.64898/2026.07.27.740904

**Authors:** Tsubasa Nishi, Izumi Kaneko, Shiroh Iwanaga, Masao Yuda

**Affiliations:** Research Institute for Microbial Diseases, Osaka University, Suita 565-0871, Japan; Department of Medicine, Mie University, Tsu 514-8507, Japan

## Abstract

Malaria sporozoites are a key developmental stage responsible for mosquito-to-mammal transmission. Sporozoites formed within oocysts on the mosquito midgut first mature into a salivary gland-invasive stage and subsequently become liver-infective upon colonizing the salivary glands. This functional switch is essential for successful parasite transmission, yet its underlying molecular mechanisms remain unclear. Here, we identify the AP2-family transcription factor AP2-Sp3 as a key regulator of this process in *Plasmodium berghei*. AP2-Sp3 is expressed in midgut sporozoites but disappears in salivary gland sporozoites. It directly binds to the *cis*-regulatory motif CATTG and represses the target genes, which include the majority of known liver infection-related genes. Disruption of *ap2-sp3* impairs development of midgut sporozoites and causes a premature shift of their transcriptome toward a salivary gland sporozoite-like state. Together, these findings demonstrate that AP2-Sp3 maintains the midgut sporozoite-specific transcriptome by suppressing the salivary gland sporozoite transcriptional program, thereby controlling the mosquito-to-mammal infectivity switch.

## Introduction

Malaria is caused by *Plasmodium* parasites, which are transmitted between humans through the bites of infected mosquitoes. A key stage for mosquito-to-human transmission is the sporozoite, a motile, crescent-shaped cell that is deposited into the dermis during blood feeding. Sporozoites subsequently migrate to the liver, differentiate into liver stage parasites, and produce merozoites, the erythrocyte-invasive form of the parasite^1–3^. Within mosquitoes, sporozoites transition through two functionally distinct states as they migrate from the midgut to the salivary glands while remaining morphologically indistinguishable. Midgut sporozoites (MgSps) are formed within oocysts on the basal lamina of the mosquito midgut and are specialized for migrating to the salivary glands^4^. In contrast, salivary gland sporozoites (SGSps) acquire infectivity for mammalian hosts^4,5^. Previous transcriptomic studies have demonstrated that transcriptional regulation is a key determinant of their distinct biological properties^6–11^. Particularly, SGSps upregulate genes associated with liver infection, encompassing those required for traversal of host cells in the dermis and liver sinusoidal layer, as well as for liver invasion and subsequent conversion to the liver stage in hepatocytes^12,13^. However, the underlying molecular mechanisms remain largely unknown.

*Plasmodium* parasites regulate stage-specific transcriptional programs throughout their complex life cycle using a limited repertoire of sequence-specific transcription factors, primarily APETALA2 (AP2) family proteins^14,15^. Among these, AP2-Sp is expressed from late oocysts to SGSps and functions as a master regulator of sporozoite development^16,17^. Its target genes span the entire mosquito-stage progression, including sporogony, egress from oocyst, salivary gland colonization, and productive invasion of hepatocytes. However, AP2-Sp alone cannot account for the distinct transcriptional patterns among these genes. Thus, the transcriptional transition from MgSp to SGSp is likely regulated through the coordinated action of multiple transcription factors.

During sporozoite development in *P. berghei*, two additional AP2 transcription factors, AP2-Sp2 and AP2-Sp3, are also essential^18^. Previous studies have shown that disruption of *ap2-sp2* results in developmental arrest before sporozoite formation, whereas *ap2-sp3* disruption does not affect sporozoite production in oocysts but causes a loss of SGSps^18,19^. Here, we show that AP2-Sp3 represses liver infection-related genes in MgSps, thereby controlling the switch in sporozoite infectivity from the salivary gland-targeting MgSp to the mammalian-infective SGSp stage.

### AP2-Sp3 is expressed in midgut and hemolymph sporozoites

AP2-Sp3 is an AP2 family transcription factor that has an AP2 domain near the center of its amino acid sequence (Extended Data Fig. 1a). BLASTP analysis detected putative orthologs of AP2-Sp3 in some apicomplexan parasites (Extended Data Fig. 1b and 1c). We elucidated its expression pattern using transgenic parasites that express GFP-tagged AP2-Sp3 (AP2-Sp3::GFP, Supplementary Fig. 1). In the mammalian host, asexual blood stages and gametocytes lacked detectable GFP signals (Extended Data Fig. 2a). Within the mosquito vector, GFP-positive oocysts emerged at 10 days post-infectious blood meal (dpi), and the proportion of GFP-positive oocysts increased over time (Fig. 1a and 1b). In addition, most GFP-positive oocysts contained sporozoites, suggesting that AP2-Sp3 expression begins during or immediately after sporogony (Fig. 1a). AP2-Sp3 expression was further observed in MgSps and over 75% of hemolymph sporozoites (HemSps), whereas the signal was completely absent in SGSps (Fig. 1a and Extended Data Fig. 2b). These results indicate that AP2-Sp3 specifically functions during MgSp-to-HemSp development.

**Fig. 1.**
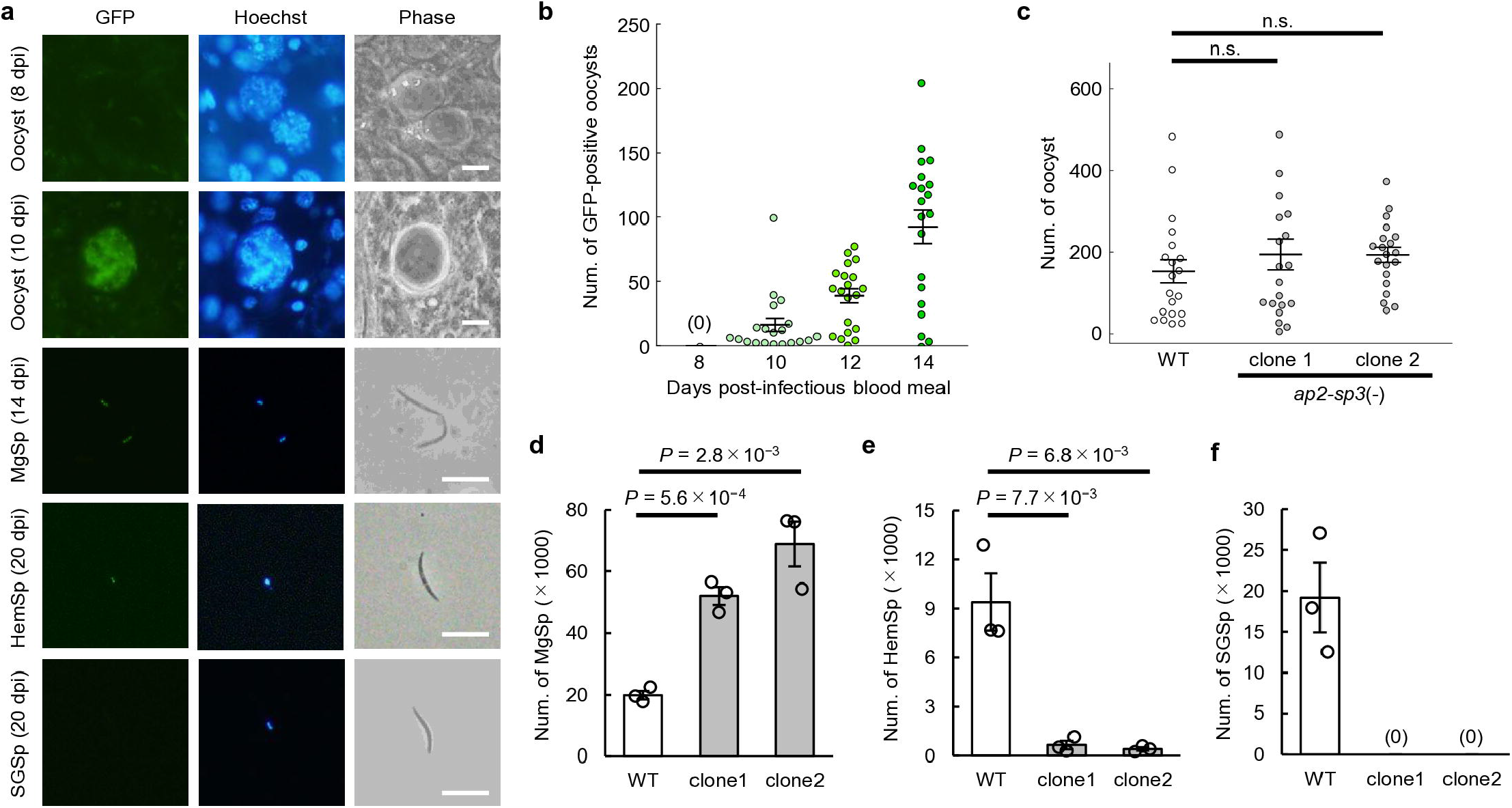
AP2-Sp3 functions during midgut-to-hemolymph sporozoite development. (a) Expression of GFP-tagged AP2-Sp3 in the AP2-Sp3::GFP parasite during mosquito-stage development. Nuclei were stained with Hoechst 33342. Scale bar = 5 μm. (MgSp: midgut sporozoite, HemSp: hemolymph sporozoite, SGSp: salivary gland sporozoite, dpi: days post-infectious blood meal) (b) The number of GFP-positive oocysts of AP2-Sp3::GFP per infected mosquito from 8 to 14 dpi (n = 20). At 8 dpi, no oocysts showed GFP signals [indicated as (0)]. (c) The number of oocysts per mosquito infected with wild-type and *ap2-sp3*(-) at 14 dpi. Lines indicate the mean values and the standard error of the mean (SEM) (n = 20). n.s. indicates *P* > 0.05 by two-tailed Student’s t-test. (d) The number of MgSps per mosquito at 14 dpi. Ten midguts were pooled for each experiment. Error bars indicate SEM from three biologically independent experiments. *P* values were calculated by two-tailed Student’s t-test. (e) The number of HemSps per mosquito at 20 dpi. Hemolymph from ten mosquitoes was pooled for each experiment. (f) The number of SGSps per mosquito at 21 dpi. Salivary glands from ten mosquitoes were pooled for each experiment. No SGSps were detected for either clone of *ap2-sp3*(-) [indicated as (0)].

### AP2-Sp3 is essential for MgSp maturation

We disrupted *ap2-sp3* using double homologous recombination [*ap2-sp3*(-), Supplementary Fig. 2]. Two independent *ap2-sp3*(-) clones were obtained and assessed for mosquito-stage development. At 14 dpi, *ap2-sp3*(-) parasites produced numbers of oocysts comparable to those of wild type (WT) (Fig. 1c); however, the number of MgSps was significantly increased (Fig. 1d). In contrast, the number of HemSps at 20 dpi was significantly lower than that in WT (Fig. 1e). Furthermore, no *ap2-sp3*(-) sporozoites were observed in salivary glands at 20 dpi (Fig. 1f). These results suggest that while *ap2-sp3*(-) forms sporozoites, their maturation within oocysts is impaired, resulting in markedly reduced egress from oocysts and a complete loss of migration to the salivary glands.

### AP2-Sp3 is required for maintaining MgSp-specific gene expression

To determine how disruption of *ap2-sp3* affects gene expression, we compared the transcriptomes of WT and *ap2-sp3*(-) MgSps at 14 dpi by bulk RNA sequencing (RNA-seq) analysis. In *ap2-sp3*(-), 311 genes were significantly upregulated [fold change > 3, Benjamini-Hochberg-adjusted *P* value (*P*^adj^) < 0.05], while 162 genes were significantly downregulated (fold change < 1/3, *P*^adj^ < 0.05) compared with WT (Fig. 2a and Supplementary Table 1). The upregulated genes included several genes associated with liver infection, such as *uis3* and *uis4*, which are known to be preferentially expressed in SGSps^20–23^ (Fig. 2a). By contrast, downregulated genes included *crmp4*, *maebl*, and *trep*, which are associated with MgSp functions, such as oocyst egress and salivary gland invasion^24–26^ (Fig. 2a). In addition, some inner membrane complex (IMC)-related genes, such as *mtip* and *phil1*, were also downregulated in *ap2-sp3*(-) (Fig. 2a).

**Fig. 2.**
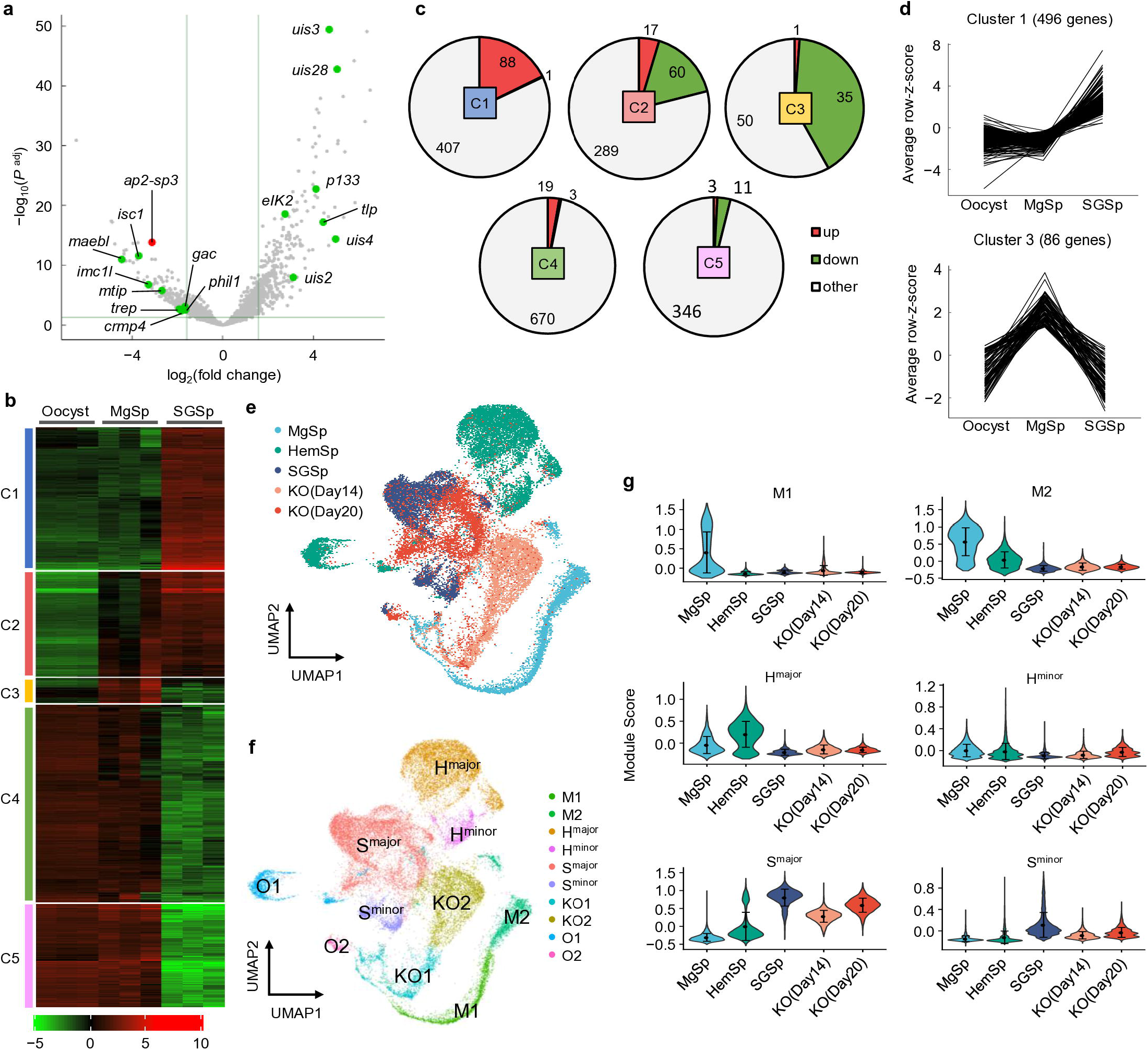
Disruption of *ap2-sp3* induces a salivary gland sporozoite-like transcriptional program in midgut sporozoites. (a) Volcano plot showing differential expression of genes in *ap2-sp3*(-). The *ap2-sp3* gene is indicated by a red dot. Green dots represent sporozoite-related genes among the differentially expressed genes. A horizontal line indicates a Benjamini-Hochberg-adjusted *P* value (*P*^adj^) of 0.05, and two vertical lines indicate fold change of 3 and 1/3. (b) Heatmap showing expression patterns of the 2,000 most variable genes during mosquito-stage development. Transcriptome datasets of oocysts [14 days post-infectious blood meal (dpi)], midgut sporozoites (MgSp) (14 dpi), and salivary gland sporozoites (SGSp) (24 dpi) were used. Genes were classified into five clusters (C1–C5) based on their expression patterns by *k*-means clustering. (c) Pie charts showing the number of genes upregulated and downregulated in *ap2-sp3*(-) for clusters C1–C5. (d) Line plots showing expression patterns of C1 and C3 genes during mosquito-stage development. (e) Uniform manifold approximation and projection (UMAP) visualization of sporozoites from wild-type (WT) and *ap2-sp3*(-). WT sporozoites were collected from the midguts (MgSp) at 14 dpi, and from the hemolymph (HemSp) and salivary glands (SGSp) at 20 dpi. For *ap2-sp3*(-), midgut sporozoites were collected at 14 [KO(Day14)] and 20 dpi [KO(Day20)]. (f) UMAP showing clusters identified by Louvain algorithm. Clusters were named based on the major cell populations they contained. O1 and O2 represent outlier clusters that did not align with the MgSp-to-SGSp developmental progress. (g) Violin plots showing distribution of cluster-specific module scores across samples. Dots and lines indicate mean values and standard deviation, respectively.

We further classified these differentially expressed genes according to their expression profiles during mosquito-stage development by *K*-means clustering. Based on our previous transcriptome datasets of oocysts and SGSps^17^ together with the current MgSp transcriptomic data, five gene clusters (C1–5) were generated from 2,000 most variable genes (Fig. 2b and Supplementary Table 2). Among these clusters, the upregulated and downregulated genes were predominantly enriched in C1 and C3, respectively (Fig. 2c and Supplementary Table 2). C1 genes were expressed at low levels in oocysts and MgSps but were highly upregulated in SGSps (Fig. 2b and 2d). In contrast, C3 genes exhibited an MgSp-specific pattern, being upregulated from oocysts to MgSps and subsequently downregulated in SGSps (Fig. 2b and 2d). Together with the phenotypic analysis, these results indicate that AP2-Sp3 plays a key role in maintaining the MgSp-specific transcriptome, which is likely essential for establishing the MgSp-specific cellular properties, such as egress from oocysts and salivary gland colonization.

### Single-cell transcriptomes of *ap2-sp3*(-) MgSps shift toward an SGSp-like pattern

Given the reciprocal upregulation of SGSp genes and downregulation of MgSp genes induced by *ap2-sp3* disruption, we hypothesized that the transcriptional profiles of *ap2-sp3*(-) sporozoites might shift toward the SGSp-like state. To examine such transcriptional transitions at the single-cell level, we performed single-cell RNA sequencing (scRNA-seq) of *ap2-sp3*(-) MgSps at 14 and 20 dpi and obtained 5,263 and 4,601 individual sporozoite transcriptomes, respectively. In addition, 4,573, 9,260, and 6,984 transcriptomes were obtained from WT MgSp (14 dpi), HemSp (20 dpi), and SGSp (20 dpi), respectively. After merging the datasets and excluding genes detected in fewer than 100 cells, expression profiles of 4,025 genes were retained for downstream analyses. A uniform manifold approximation and projection (UMAP) embedding was then generated based on principal component analysis (PCA) (Fig. 2e), and clustering analysis identified multiple clusters among all cells (Fig. 2f). Transcriptomes of WT MgSps appeared to progress from M1 to M2, representing MgSp development within oocysts, and subsequently to HemSp clusters (Fig. 2e and 2f). HemSps and SGSps were mainly separated into two clusters: one included the majority of the population (H^major^ and S^major^), and the other represented a minor population (H^minor^ and S^minor^) (Fig. 2e and 2f). On the UMAP, *ap2-sp3*(-) MgSps at 14 dpi followed a transcriptomic progression distinct from that of WT (from clusters KO1 to KO2) and instead shifted toward SGSp clusters (Fig. 2e and 2f). Furthermore, most 20 dpi cells clustered within S^major^ (Fig. 2e and 2f). In module score analysis using cluster-specific marker genes, *ap2-sp3*(-) MgSps exhibited the highest score for S^major^ among the other clusters (Fig. 2g). These results are consistent with the bulk RNA-seq results, which demonstrated upregulation of SGSp-specific genes and downregulation of MgSp-specific genes in *ap2-sp3*(-) MgSps. Together, these results indicate that *ap2-sp3*(-) sporozoites prematurely activated the SGSp-specific transcriptional program while failing to properly establish the MgSp-specific transcriptional program.

### The SGSp-like transcriptional state of *ap2-sp3*(-) does not confer mammalian infectivity

Given the SGSp-like transcriptome, we asked whether *ap2-sp3*(-) MgSps had acquired infectivity for mammalian hosts. To assess their infectivity, we collected MgSps of WT and *ap2-sp3*(-) at 14 dpi and inoculated 50,000 sporozoites per rat. In WT, blood-stage parasites were observed in six of eight rats injected with MgSps, with a mean prepatent period of 5.2 days (Extended data Table 1). In contrast, *ap2-sp3*(-) parasites were detected in only one of eight rats with a prepatent period of 6.0 days (Extended data Table 1). These results indicate that acquisition of mammalian infectivity requires the prior completion of MgSp maturation, and that a mere transcriptomic shift is insufficient.

### Direct binding to CATTG via the AP2 domain is essential for AP2-Sp3 function

To understand how the transcriptional transition from MgSp to SGSp was induced by *ap2-sp3* disruption, we next performed chromatin immunoprecipitation followed by sequencing (ChIP-seq) analysis using AP2-Sp3::GFP at 14 dpi (Extended Data Fig. 3a). ChIP-seq was performed in duplicate, yielding 1,428 and 1,528 peaks, respectively, with 1,183 peaks overlapping (83% of peaks in experiment 1) (Extended Data Fig. 3b and 3c, and Supplementary Tables 3a and 3b). We then performed motif enrichment analysis using Fisher’s exact test and identified two motifs enriched within these common peak regions, CATTG (*P* = 4.8 × 10^−162^) and TGCATG (*P* = 3.1 × 10^−160^) (Supplementary Table 3c and 3d). These motifs were found within 300 bp of the peak summit of 856 and 532 peaks, respectively, and were highly enriched around the peak summits (Fig. 3a and 3b).

**Fig. 3.**
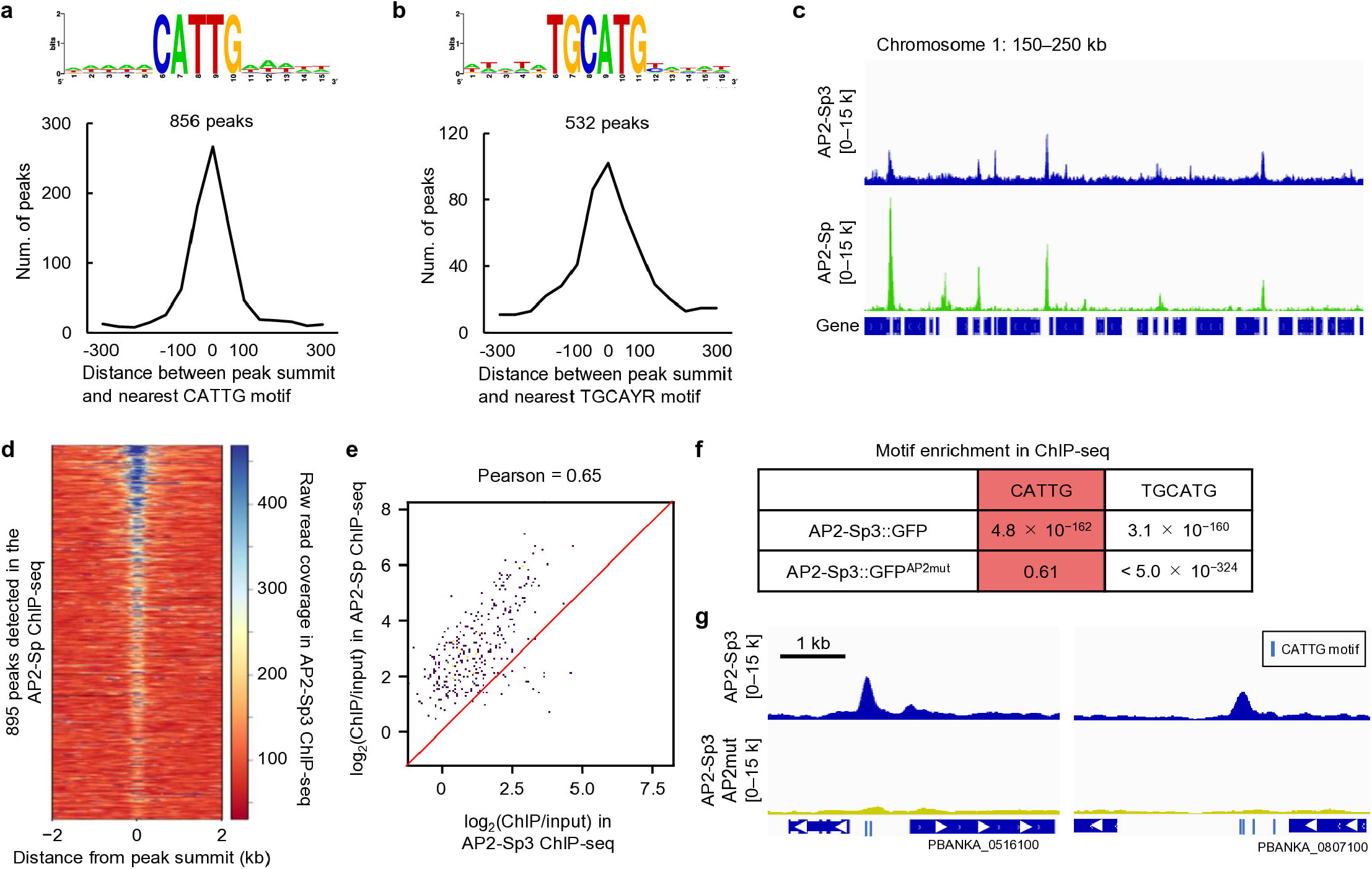
AP2-Sp3 targets sporozoite-related genes. (a, b) Motifs enriched within 50 bp from the AP2-Sp3 peak summits: (a) CATTG and (b) TGCATG. The logo was created using WebLogo (https://weblogo.berkeley.edu/logo.cgi). The line plot at the bottom shows the number of peaks along the distance between the peak summits and the nearest motif. The total number of peaks containing CATTG and TGCATG within 300 bp of the summit is shown at the top of each plot. (c) Integrative Genomics Viewer (IGV) images of AP2-Sp3 and AP2-Sp ChIP-seq data for a region of chromosome 1. Histograms show the raw read coverage of ChIP data normalized by library size (bin size = 10 bp). Scales are indicated in square brackets. (d) Heatmap showing raw read coverage of AP2-Sp3 ChIP data at the AP2-Sp peaks. AP2-Sp peak regions are aligned in the ascending order of their *q* values. (e) Scatter plot comparing log_2_(ChIP/input) of AP2-Sp3 and AP2-Sp ChIP-seq at AP2-Sp peak regions. A red line indicates y = x. (f) Enrichment of CATTG and TGCATG motifs in the AP2-Sp3::GFP and AP2-Sp3::GFP^AP2mut^ ChIP-seq. *P* values were calculated by Fisher’s exact test. (g) IGV images for the peaks depleted in AP2-Sp3::GFP^AP2mut^ ChIP-seq. Blue bars indicate positions of CATTG motifs.

TGCATG is the major binding motif of AP2-Sp, a master transcription factor for sporozoite development. Comparison of ChIP-seq analyses of AP2-Sp3 and AP2-Sp revealed that besides CATTG-associated peaks, AP2-Sp3 ChIP-seq exhibited peak patterns centered around AP2-Sp binding sites (Fig. 3c and 3d). ChIP/input signals at AP2-Sp peak regions were moderately correlated between the two datasets (Pearson’s r = 0.65), and the log_2_(ChIP/input) values at these sites were generally lower in the AP2-Sp3 ChIP-seq than in the AP2-Sp ChIP-seq (Fig. 3e). These results suggest that the enrichment of TGCATG in the AP2-Sp3 ChIP-seq may be derived from indirect recruitment of AP2-Sp3 to AP2-Sp-bound loci.

To determine whether CATTG is the direct binding motif of AP2-Sp3, we performed ChIP-seq using an AP2-Sp3 mutant (AP2-Sp3::GFP^AP2mut^), carrying alanine substitutions at two apicomplexan-conserved residues within the AP2 domain (W904A and R918A) to specifically disrupt its DNA binding ability (Extended Data Fig. 1c and 4a). This analysis detected enrichment of TGCATG around the peak summits (*P* < 5.0 × 10^−324^, the smallest positive value representable in R), whereas CATTG was no longer enriched (*P* = 0.61) (Fig. 3f and Supplementary Tables 4a–c). Consistently, the peaks associated with CATTG, but not with the AP2-Sp binding motifs, were lost in the AP2-Sp3::GFP^AP2mut^ ChIP-seq (Fig. 3g). These results indicate that AP2-Sp3 directly recognizes CATTG motifs via its AP2 domain. Notably, AP2-Sp3::GFP^AP2mut^ parasites phenocopied *ap2-sp3*(-) (Extended Data Fig. 4b, 4c, 4d, and 4e). These results demonstrate that direct binding to CATTG motifs, rather than association with AP2-Sp-bound loci, is essential for AP2-Sp3 function.

**Fig. 4.**
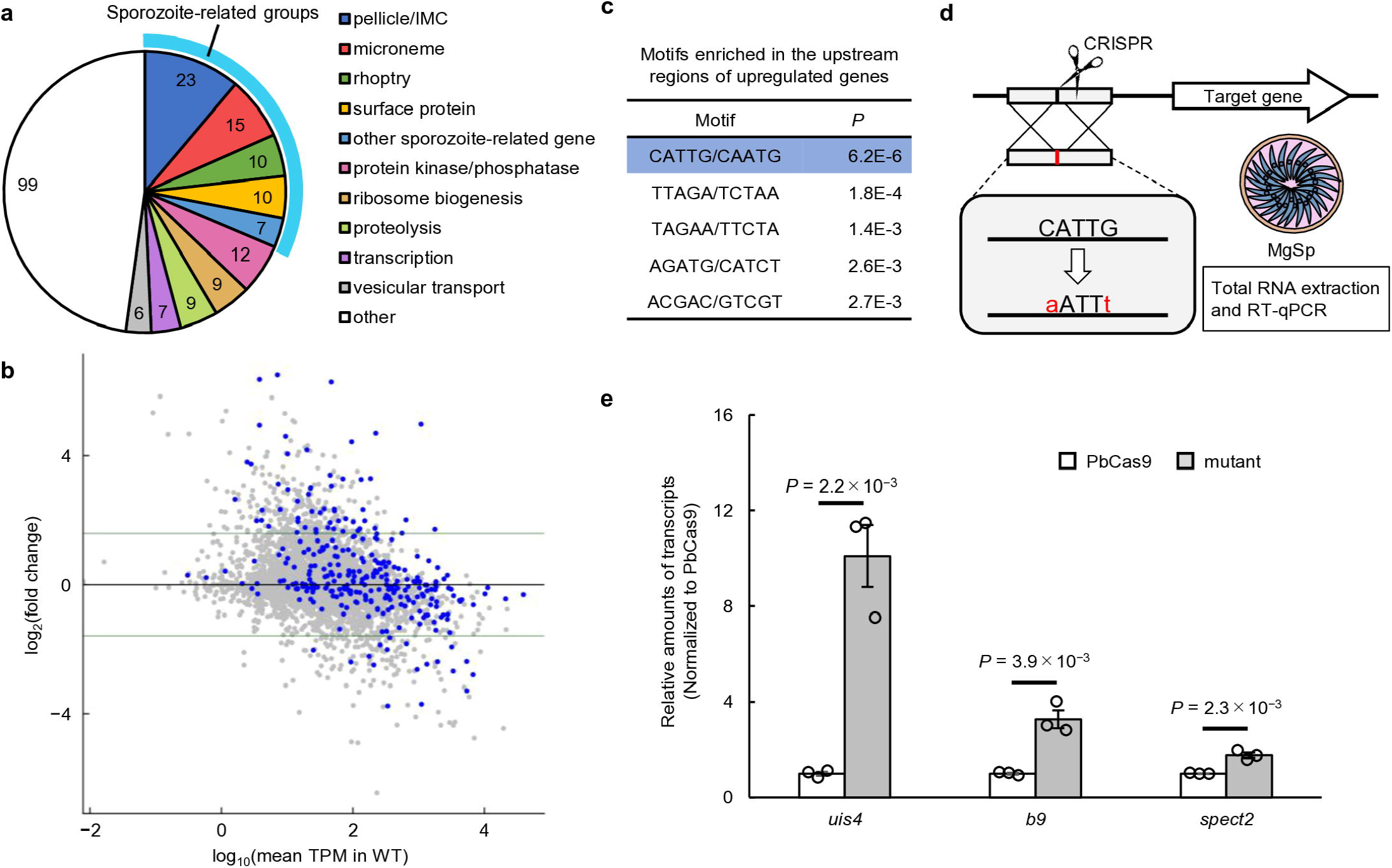
AP2-Sp3 functions as a transcriptional repressor. (a) Classification of AP2-Sp3 target genes. Groups that are associated with sporozoite structure and function are indicated by a blue line. Genes with unknown functions are excluded from the classification. (b) MA plot showing differential expression of genes between midgut sporozoites of wild-type and *ap2-sp3*(-) at 14 days post-infectious blood meal (dpi). Blue dots represent AP2-Sp3 target genes. Three horizontal lines indicate fold change of 3, 1, and 1/3. (c) Five-base pair motifs enriched in the upstream regions (200–1000 bp from start codons) of genes significantly upregulated in *ap2-sp3*(-). (d) Schematic illustration of reporter experiments. Mutations were introduced into AP2-Sp3-binding motifs (CATTG to aATTt) on endogenous promoters of target genes using the CRISPR/Cas9 system. Gene expression levels were quantified by reverse-transcription quantitative PCR (RT-qPCR) analysis. (e) RT-qPCR analysis of AP2-Sp3 target gene expression in midgut sporozoites (14 dpi) following introduction of mutations into the AP2-Sp3-binding motifs in their upstream regions. Relative transcript levels normalized to *maebl* were further normalized to the values for PbCas9. Error bars indicate the standard error of the mean from three biologically independent experiments. *P* values were calculated by two-tailed Student’s t-test.

### AP2-Sp3 acts as a transcriptional repressor

To investigate the transcriptional regulatory function of AP2-Sp3, we explored its target genes using ChIP-seq peaks containing CATTG motifs (Fig. 3b). Defining genes with a peak within 1200 bp of their start codon as targets, we identified 296 AP2-Sp3 target genes (Supplementary Table 5a). Among these, “pellicle/IMC” and “microneme” genes constituted the largest functional categories (Fig. 4a and Extended Data Fig. 5). The targets also included some rhoptry and sporozoite surface protein genes (Fig. 4a). Consistently, gene ontology analysis detected enrichment for terms related to these structures, such as “apical complex,” “movement in host,” “microneme,” and “rhoptry” (Supplementary Table 5b). Targets in “other sporozoite-related gene” included genes important for liver infection, such as *uis3* and *uis4*, as well as *puf2*, which is involved in translational repression and maintenance of liver-stage gene transcripts in SGSps^27^. (Fig. 4a and Extended Data Fig. 5). Furthermore, although excluded from the target list due to the 1200-bp threshold, another post-transcriptional regulator gene, *eIK2*^23^, harbored AP2-Sp3 ChIP-seq peaks in its upstream region (Extended Data Fig. 5).

We then examined the expression changes of these targets between WT and *ap2-sp3*(-) using the transcriptomic datasets of MgSps at 14 dpi (Supplementary Table 1). Overall, the mean log_2_(fold change) value of all target genes was significantly higher than that of non-target genes (*P* = 2.0 × 10^−10^ by two-tailed Student’s t-test) (Fig. 4b and Supplementary Table 1). In addition, the significantly upregulated genes contained 55 AP2-Sp3 target genes, showing a significant overlap with a *P* value of 4.6 × 10^−13^ by Fisher’s exact test (Fig. 4b and Supplementary Table 1). In the upstream regions (200–1000 bp from the start codon) of genes upregulated in *ap2-sp3*(-), the CATTG motif was most significantly enriched with a *P* value of 6.2 × 10^−6^ by Fisher’s exact test (Fig. 4c). These results indicate that AP2-Sp3 functions as a transcriptional repressor during MgSp development.

### CATTG functions as a *cis*-regulatory element for transcriptional repression

To determine whether the AP2-Sp3-binding motif functions as a *cis*-regulatory element mediating transcriptional repression, we introduced mutations into the motifs in the endogenous promoters of AP2-Sp3 targets using a Cas9-expressing parasite line, PbCas9^28^ (Fig. 4d). We selected three target genes, *uis4*, *b9*, and *spect2*, for the assays. Transcriptional activities of their promoters were assessed by RT-qPCR analysis using MgSps at 14 dpi, with or without mutations altering CATTG to aATTt (Fig. 4d and Extended Data Fig. 6). The analysis showed that the amounts of target transcripts relative to that of *maebl* (internal control selected due to its high expression in MgSps) significantly increased after introducing the mutations (*uis4*, 10.1-fold; *b9*, 3.3-fold; *spect2*, 1.8-fold) (Fig. 4e). This result confirms that the AP2-Sp3-binding motifs mediate transcriptional repression of target genes during MgSp development.

### AP2-Sp3 establishes MgSp-specific transcriptome by repressing two distinct gene sets

In *Plasmodium*, transcriptional repressors play multiple regulatory roles*^29–31^*. Although AP2-Sp3 represses premature activation of the SGSp-specific transcriptional program, the target genes also included many genes that are not SGSp-specific. Thus, to gain a deeper understanding of the roles of AP2-Sp3, we performed hierarchical clustering of its targets to characterize them based on their expression patterns during mosquito-stage development. This analysis classified the targets into two groups with distinct expression patterns (Fig. 5a and Supplementary Table 5a). Group 1 exhibited low expression levels from oocysts through MgSps, followed by strong induction in SGSps (Fig. 5a). This group encompassed most of the known liver infection-related genes^3,22,32–37^ (red in Fig. 5b). Furthermore, the majority of targets upregulated in *ap2-sp3*(-) were included in group 1 (Supplementary Table 5a). Group 2 genes were highly expressed in oocysts and downregulated thereafter toward SGSp (Fig. 5a). This group included several genes related to sporozoite morphogenesis and motility^38–44^ (green in Fig. 5b). In the differential expression analysis, only a limited number of group 2 targets were significantly upregulated in *ap2-sp3*(-) (Supplementary Table 5a), presumably because their high baseline expression in MgSps moderated the magnitude of increase. Collectively, these two contrasting expression patterns among the targets suggest that AP2-Sp3 fulfills two principal regulatory roles in MgSps: preventing premature activation of SGSp-specific genes, and downregulating genes that are activated for sporogony as MgSp development proceeds (Fig. 5c).

**Fig. 5.**
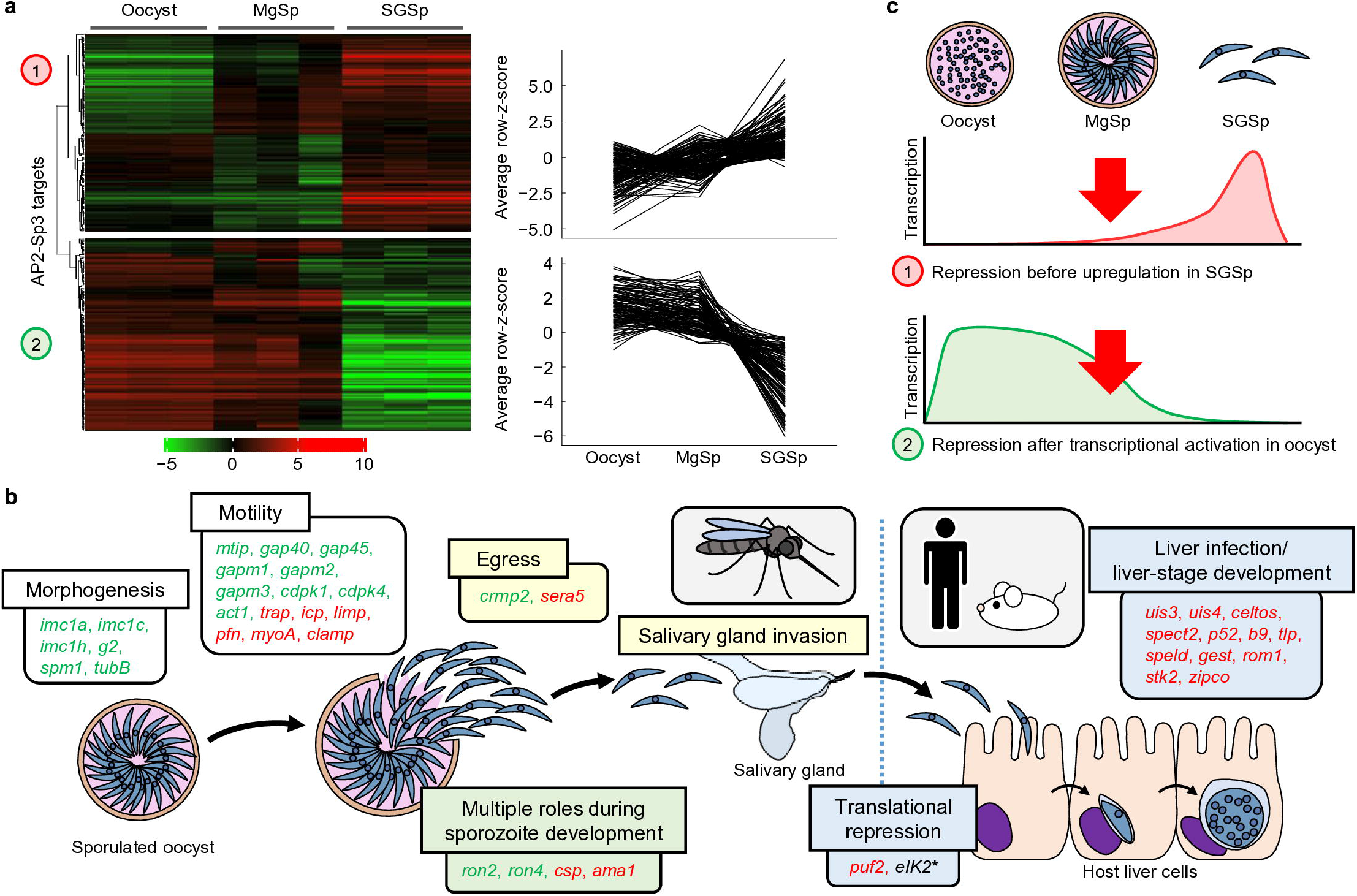
AP2-Sp3 plays two major roles in establishing the midgut sporozoite-specific transcriptional profiles. (a) Hierarchical clustering of AP2-Sp3 target genes according to their expression patterns in the transcriptome datasets of oocysts (14 dpi), midgut sporozoites (MgSps) (14 dpi), and salivary gland sporozoites (SGSps) (21 dpi). Targets are classified into two major groups. Line plots show the expression patterns of each gene in groups 1 and 2. (b) Schematic illustration of *P. berghei* sporozoite development. AP2-Sp3 targets associated with sporozoite cellular properties are shown. Targets belonging to groups 1 and 2 are indicated in red and green, respectively. Group 1 mainly includes those associated with liver infection and liver-stage development, whereas group 2 includes genes associated with sporozoite morphogenesis and motility that are expressed during sporogony. (c) Schematic representation of expression patterns of targets in groups 1 and 2 during sporozoite development.

## Discussion

*Plasmodium* sporozoites undergo a marked change in infectivity during development within the mosquito host^4,45^. This infectivity switch has been thought to be achieved through the regulation of genes involved in liver infection, but the underlying molecular mechanisms remained unclear. In this study, we identified AP2-Sp3 as a key regulator of this MgSp-to-SGSp transcriptional transition. AP2-Sp3 functions as a transcriptional repressor in MgSps and represses SGSp-specific genes, which encompass the majority of genes essential for liver infection and intrahepatic development. Disruption of *ap2-sp3* relieved this repression and converted the MgSp transcriptome into an SGSp-like state. These results demonstrate that expression of liver infection-related genes is strictly regulated by AP2-Sp3 and establish the AP2-Sp3-mediated repression as the molecular basis of the infectivity switch in sporozoites.

This study also demonstrated that AP2-Sp3 plays a decisive role in the maturation of MgSps within the oocyst. AP2-Sp3 safeguards against the premature activation of the SGSp-specific transcriptional program and also represses genes highly expressed during sporogony as suggested by the target gene analysis. These transcriptional repressor activities likely contribute to the establishment of the MgSp-specific transcriptome. Consequently, in *ap2-sp3* knockout parasites, the expression of MgSp-specific genes was downregulated, which impaired normal MgSp development and caused severe defects in oocyst egress and salivary gland colonization.

Another key finding of this study is that an SGSp-like transcriptome of *ap2-sp3* knockout parasites does not confer liver infectivity. Although *ap2-sp3* knockout MgSps prematurely activated the SGSp-specific transcriptional program, they exhibited markedly reduced liver infectivity. This result indicates that the formation of fully liver-infective sporozoites requires not only activation of the SGSp-specific transcriptional program but also completion of the MgSp maturation program. It further suggests that AP2-Sp3-mediated transcriptional repression must be maintained throughout MgSp development and relieved only after MgSp maturation is complete to enable proper SGSp formation.

An important unanswered question is how AP2-Sp3 disappears during sporozoite development. Because AP2-Sp3 is expressed in MgSps and most HemSps but is completely absent in SGSps, its disappearance is likely triggered by extrinsic cues associated with salivary gland invasion. One plausible mechanism is that the recognition of the salivary gland by adhesion molecules, such as MAEBL, or the initiation of salivary gland invasion, activates a signaling pathway that promotes AP2-Sp3 degradation. This model suggests that host–parasite interactions can directly regulate developmental transitions in *Plasmodium*. Elucidating the molecular mechanisms linking salivary gland invasion to AP2-Sp3 degradation will therefore be an important direction for future studies.

Recently, techniques for *in vitro* sporozoite production have advanced, raising expectations for their application as live-attenuated vaccines. However, the infectivity of these *in vitro*-generated sporozoites remains substantially lower than that of mosquito-derived sporozoites^46^. This study demonstrates that the appropriate developmental stage transition, orchestrated by AP2-Sp3, is indispensable for sporozoites to acquire full infectivity. In the future, elucidating the signaling mechanisms that induce AP2-Sp3 degradation and artificially replicating them *in vitro* could provide a breakthrough in producing highly liver-infective *in vitro*-generated sporozoites.

In summary, this study demonstrates that the infectivity switch from MgSps to SGSps is initiated not by the emergence of a novel transcriptional activator, but by the disappearance of the transcriptional repressor AP2-Sp3. While previous studies have reported the critical roles of transcriptional repressors in stage-specific gene expression at multiple developmental stages of the *Plasmodium* life cycle^29–31^, our findings clearly establish their importance in sporozoite development as well. By working in concert with transcriptional activators that similarly control an extensive array of target genes, these wide-ranging transcriptional repressors likely enable the generation of the diverse transcriptomes required for each life cycle stage using a limited repertoire of transcription factors.

## Methods

### Ethical statement

All experiments involving mice were performed according to the recommendations in the Guide for the Care and Use of Laboratory Animals of the National Institutes of Health in order to minimize animal suffering. These experiments were approved by the Animal Research Ethics Committee of Mie University (permit number 23–29) and Animal Experiments Committee of Osaka University (permit number R07-11-0).

### Parasite preparation

Parasites were inoculated into ddY mice for generating transgenic parasites. For mosquito infection experiments, parasites were propagated in BALB/cCrSlc mice. Sporozoite infectivity assays were performed using Wistar rats.

Female *Anopheles stephensi* mosquitoes, 5–7 days post-emergence, were used for *P. berghei* infection. Prior to the infectious blood meal, mosquitoes were starved for 6–12 h at 20 °C to enhance feeding efficiency and acclimate them to this temperature. Subsequently, the mosquitoes were allowed to feed on *P. berghei*-infected mice at 20 °C. After feeding, engorged mosquitoes were collected in a new cage and were maintained at 20 °C with access to 5% D-fructose solution.

To harvest midgut and salivary gland sporozoites, infected mosquitoes were dissected, and midguts or salivary glands were homogenized in PBS using a pestle. Sporozoites were then separated from tissue debris by centrifugation at 500 × g for 2 min. For collection of hemolymph sporozoites, a small cut was made in the mosquito abdomen using a fine needle. Subsequently, 10 μL of MEM was injected into the thorax using a microsyringe fitted with a thin, custom-made glass needle, and hemolymph expelled from the abdomen incision was collected.

### Plasmid construction and generation of transgenic parasites

The *ap2-sp3*(-) parasite line was generated from WT *P. berghei* ANKA strain by the conventional homologous recombination method. The targeting construct, which was composed of two homologous regions around the *ap2-sp3* locus flanking a *hdhfr* expression cassette, was produced by overlap PCR. All other transgenic parasite lines were derived from the PbCas9 parasite line, which constitutively expresses Cas9 nuclease^28^. Donor DNAs were constructed by overlap PCR and cloned into the *Xho*I and *Bam*HI sites of pBluescript KS (+) using In-Fusion cloning (Takara). Donor DNAs were then amplified from the plasmid by PCR. Target sequences of single guide RNA (sgRNA) were designed using CHOPCHOP (https://chopchop.cbu.uib.no/), and sgRNA templates were constructed by annealing DNA oligos and then were cloned into the previously reported sgRNA vector using the DNA Ligation Kit (Takara)^28^.

Transfection was performed as previously described. Briefly, parasites were cultured in RPMI 1640 medium supplemented with 25% fetal calf serum and penicillin/streptomycin for 16 h at 37 °C under 5% CO_2_ and 10% O_2_ conditions. Schizonts produced in the cultures were harvested by density gradient centrifugation and were then transfected with DNA constructs using Amaxa Basic Parasite Nucleofector Kit 2 (LONZA), followed by inoculation into mice. Transfectants were selected by treatment of mice with 70 μg/mL pyrimethamine in their drinking water. Recombination was confirmed by PCR and/or Sanger sequencing, and clonal parasites were obtained by limiting dilution. All primers used in this study were listed in Supplementary Table 6.

### Fluorescence analysis

GFP fluorescence was analyzed using an Olympus BX51 microscope with Olympus DP74 camera. To visualize nuclei, midguts and salivary glands were suspended in PBS containing 2 ng/mL Hoechst 33342 and incubated for 10 min at room temperature. For hemolymph sporozoites, Hoechst 33342 solution was added to extracted hemolymph to reach a final concentration of 2 ng/mL, and the mixture was incubated for 10 min at room temperature. Blood-stage parasite nuclei were stained with 1 ng/mL Hoechst 33342 for 10 min at 37 °C.

### Phenotype analysis of mosquito-stage parasites

At 14 dpi, infected mosquitoes were dissected, and the number of oocysts per midgut was counted under a phase-contrast microscope. To quantify sporozoites, midguts (14 dpi), hemolymph (20 dpi), or salivary glands (20 dpi) from ten mosquitoes were pooled for each replicate. Sporozoites in each sample were counted using a hemocytometer. Three biologically independent experiments were conducted for sporozoite quantification.

For sporozoite infectivity assays, 50,000 midgut sporozoites or 1,000 salivary gland sporozoites were injected intravenously into Wistar rats. Prepatent days were determined by daily Giemsa staining of peripheral blood collected from the tail vein.

### Single-cell RNA-seq and sequence data analysis

Sporozoites were harvested and pooled from 15 mosquitoes as described for parasite preparation. Cells were passed through a 20-μm CellTrics filter (Sysmex) to remove tissue debris, and the buffer was replaced with PBS containing 0.04% BSA. Sporozoites were counted using a hemocytometer, and the cell concentration was adjusted to 800–1200 cells/μL. Single-cell RNA-seq libraries were generated using the Chromium GEM-X Single Cell 5’ Kit v3 (10x Genomics) according to the manufacturer’s instructions. Libraries were sequenced on a NovaSeq X Plus platform (Illumina) with 150-bp paired-end reads, generating approximately 200 million reads per sample.

Raw sequencing reads were processed with Cell Ranger v9.0.1 (10x Genomics) count pipeline. Reference genome sequence and gene annotations were obtained from PlasmoDB release 68^47^. Because untranslated regions are not annotated in *P. berghei*, the 5’ end of each gene was extended by 300 bp to improve read assignment efficiency. Multigene families (*pir* and *fam*) were excluded from downstream analyses. Raw count matrices were imported into R using the Read10X function in Seurat v5.4.0. For each sample, cells with more than 500 total RNA counts (nCount_RNA) and more than 100 detected genes (nFeature_RNA) were retained. Doublets were identified using scDblFinder and removed prior to the following analyses. Filtered Seurat objects from all samples were merged into a single object, and genes detected in at least 100 cells across all samples were retained. The filtered dataset was normalized and variance-stabilized using SCTransform. Dimensionality reduction was performed by PCA, followed by visualization using UMAP based on the top 20 principal components. Cell clustering was performed using FindNeighbors and FindClusters in Seurat with a clustering resolution of 0.1.

Cluster marker genes were identified using the FindAllMarkers function in Seurat, setting a minimum expression frequency to 25% and a log_2_(fold change) threshold to 0.25, after SCT assay preparation with PrepSCTFindMarkers. For each cluster, the top 10 marker genes ranked by average log_2_(fold change) were selected to generate cluster-specific gene signatures. Module scores for each signature were calculated using the AddModuleScore function in Seurat. Parameters for all programs were set as default unless otherwise specified.

### RNA-seq and sequence data analysis

Midgut sporozoites were collected from 100 mosquitoes per sample at 14 dpi. The sporozoite homogenate was further passed through a DEAE ion-exchange column to remove tissue debris. The purified sporozoites were then lysed in Isogen II (NIPPON GENE). Total RNA was extracted from the Isogen II lysates according to the manufacturer’s instructions. RNA-seq libraries were then prepared using KAPA mRNA HyperPrep Kit (Kapa Biosystems) and were sequenced by the MGI DNBSEQ-G400. Three biologically independent experiments were performed for each genotype. The sequence data were mapped onto the reference *P. berghei* ANKA genome by HISAT2, with the maximum intron length set to 1000. Mapped read counts for each gene were calculated using featureCounts. The differential expression analysis between WT and *ap2-sp3*(-) was performed using DESeq2, applying apeglm shrinkage to evaluate the effect sizes without arbitrarily filtering out genes with low counts. Multigene families (*pir* and *fam*) were excluded from the differential expression analysis. Parameters for all programs were set as default unless otherwise specified. Gene clustering analysis was performed using the iDEP 2.0 platform with the number of clusters set to five. The 2,000 most variable genes were classified into these clusters.

### ChIP-seq and sequencing data analysis

ChIP-seq was performed as previously described for AP2-Sp ChIP-seq^17^. Briefly, 100 infected mosquitoes were dissected, and midguts were collected in PBS containing 1% formalin at room temperature. After collection, the suspension was further incubated for 1 h at 30 °C. The fixed midguts were washed once with PBS, suspended in SDS lysis buffer (50 mM Tris-HCl, 1% SDS, 10 mM EDTA), and then sonicated using a Bioruptor sonicator (Cosmo Bio). Using the sonicated lysate, chromatin immunoprecipitation was performed with anti-GFP polyclonal antibodies (ab290, Abcam) conjugated to Dynabeads Protein A (Invitrogen). Following overnight incubation with rotation at 4 °C, the beads were washed five times with low-salt wash buffer (20 mM Tris-HCl, 0.1% SDS, 2 mM EDTA, 1% Triton X-100, 150 mM NaCl) and three times with high-salt wash buffer (20 mM Tris-HCl, 0.1% SDS, 2 mM EDTA, 1% Triton X-100, 500 mM NaCl). The beads were then resuspended in extraction buffer (10 mM Tris-HCl, 1% SDS, 5 mM EDTA, 300 mM NaCl) and were incubated with rotation for 15 min at room temperature to extract the immunoprecipitated chromatin. The extract was incubated for 6 h at 65 °C and treated with RNase A (50 μg/mL) for 1 h at 37 °C followed by Proteinase K treatment (500 μg/mL) for 2 h at 55 °C. DNA fragments were purified from the immunoprecipitated chromatin by phenol/chloroform extraction, and libraries were prepared using KAPA HyperPrep Kit (Kapa Biosystems). Next-generation sequencing (NGS) was performed using the MGI DNBSEQ-G400. Small aliquots of lysate before immunoprecipitation were used for obtaining input sequence data. Two biologically independent ChIP-seq experiments were performed for each parasite line, except for the ChIP-seq analysis using AP2-Sp3::GFP^AP2mut^, which was only conducted for motif enrichment analysis.

Sequence reads were mapped to the reference genome sequence of *P. berghei* (v3.0 downloaded from PlasmoDB) by Bowtie 2. Reads mapped to more than two genomic locations were removed from the mapping data using the grep utility, and the mapped sequence data were converted from SAM to sorted BAM using samtools sort. Peaks were called by macs2 callpeak using input sequence data as a control (fold enrichment > 2.0, *q* value < 0.01). Peaks were compared between biological replicates using IDR1D (https://idr2d.mit.edu/)^48^ (max gap = 100) to evaluate the reproducibility. Pearson’s correlation coefficient between the AP2-Sp3 and AP2-Sp ChIP-seq datasets was calculated using multiBigwigSummary bins (-bs 1000). Common peaks in biological replicates were defined as those with a distance less than 150 bp between their peak summits and used for the following analyses. Enrichment of motifs within 50 bp from peak summits relative to the rest of the genome sequence was analyzed using Fisher’s exact test. For the motif enrichment analysis, peaks located within 1 kb of the end of each chromosome were excluded to avoid detection of the telomeric repeat sequence, TTYAGGG (Y = T or C). Genes that have peaks within their 1200-bp upstream intergenic regions were identified as target genes. Gene ontology analysis was performed using GSEABase and GOstats programs. All enriched terms from three aspects, Molecular Function, Cellular Component, and Biological Process, were ranked along their *p*-values. Hierarchical clustering analysis of target genes was performed using the iDEP 2.0 platform. Parameters for all programs were set as default unless otherwise specified.

### RT-qPCR analysis

Total RNA of midgut sporozoites was harvested as described for RNA-seq analysis, except that only 20 midguts were collected per sample. From the total RNA, cDNA was synthesized using PrimeScript RT reagent Kit with gDNA Eraser (Takara). The amounts of target gene cDNA were quantified using TB Green Fast qPCR Mix (Takara) and Thermal Cycler Dice Real Time System II (Takara), setting amplification cycles as 40 cycles. Three biologically independent samples were prepared for each experiment. All primers used were listed in Supplementary Table 6.

## Statistical analysis

Statistical significance was determined using two-sided Student’s t-tests in Microsoft Excel or Fisher’s exact tests in R. For differential expression analysis, the Benjamini-Hochberg-adjusted *P*-values were calculated using DESeq2. Sample sizes were chosen based on common practice in similar studies. No experiments were randomized or blinded because the compared samples were genetically distinct (for example, wild-type and knockout lines) or represented different developmental stages (for example, midgut and salivary gland sporozoites). All values are presented as mean ± s.e.

## Data availability

All FASTQ files for scRNA-seq, RNA-seq, and ChIP-seq experiments are available from the Gene Expression Omnibus database (accession numbers GSE333890, GSE334238, and GSE333780).

## Acknowledgements

We thank the NGS core facility at the Research Institute for Microbial Diseases of Osaka University for assistance with 10x Genomics single-cell RNA-seq library preparation and sequencing.

---

**Extended Data Fig 1.**
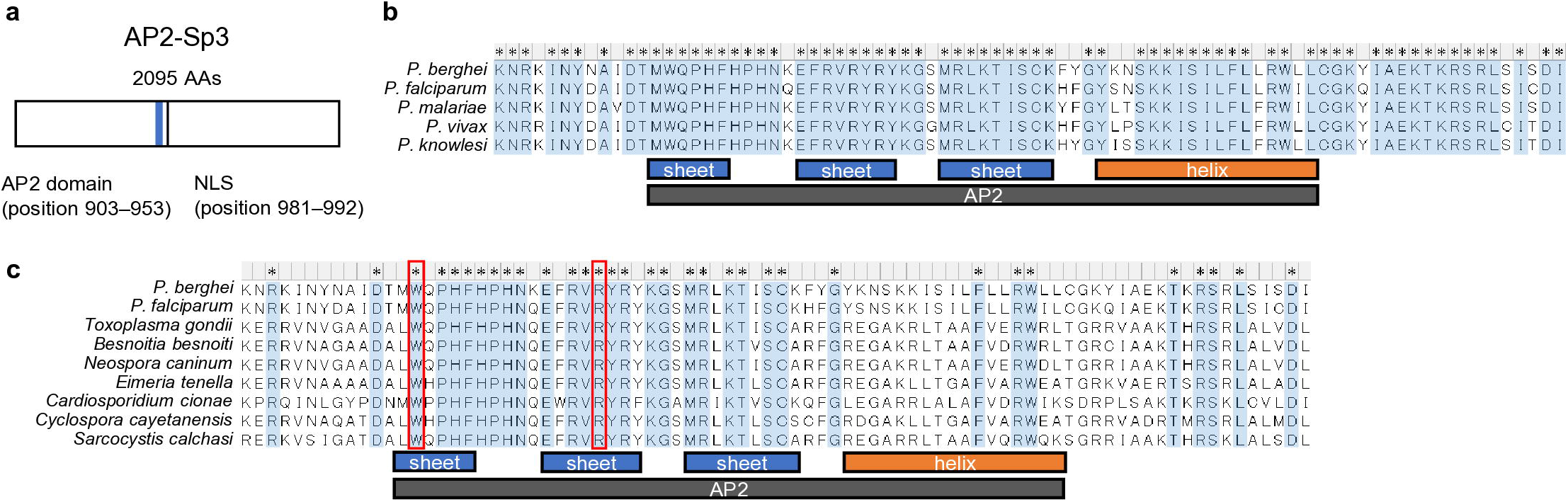
AP2-Sp3 is an apicomplexan-conserved AP2 family member. (a) Schematic illustration of AP2-Sp3. Region of the AP2 domain is indicated by blue. Nuclear localization signal (NLS) was predicted using cNLS Mapper (http://nls-mapper.iab.keio.ac.jp/cgi-bin/NLS_Mapper_form.cgi) and is indicated by black. (b) Alignment of amino acid sequences for the AP2 domain of AP2-Sp3 and its orthologs in *Plasmodium* by the ClustalW program in Mega X (*P*. *berghei*, PBANKA_0909600; *P*. *falciparum*, PF3D7_1139300; *P. malariae*, PmUG01_09048500; *P. vivax*, PVP01_0940100; *P*. *knowlesi*, PKNH_0937300; *P. gallinaceum*, PGAL8A_00366800). Amino acids that are conserved in all orthologs are highlighted by blue and an asterisk. Region of an AP2 domain and α-helix and β-sheets within it are indicated by boxes below sequence alignment. (c) Alignment of amino acid sequences for the AP2 domain of AP2-Sp3 and its putative orthologs in *Apicomplexa* (*Toxoplasma gondii*, XP_018638208; *Besnoitia besnoiti*, XP_029215173; *Neospora caninum*, XP_003880306; *Eimeria tenella*, XP_013227991; *Cardiosporidium cionae*, KAF8821986; *Cyclospora cayetanensis*, XP_026192272; *Sarcocystis calchasi*, CAL7866311).

**Extended Data Fig 2.**
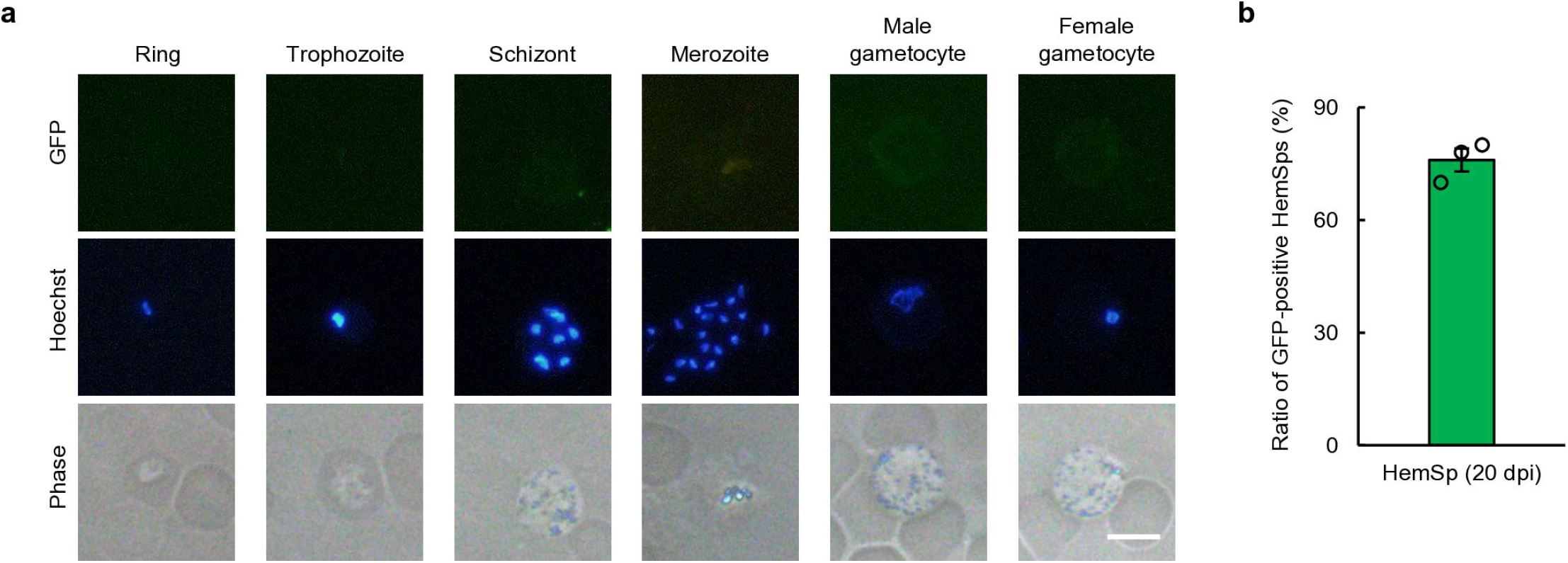
AP2-Sp3 is not expressed in blood stage parasites. (a) Expression of AP2-Sp3 in the AP2-Sp3::GFP parasite during blood stage. Nuclei were stained with Hoechst 33342. Scale bar = 5 μm. (b) Ratio of GFP-positive hemolymph sporozoites (HemSps) in AP2-Sp3::GFP at 20 days post-infectious blood meal (dpi). Error bars indicate the standard error of the mean (SEM) from three biologically independent experiments. For each experiment, hemolymph from ten mosquitoes were pooled, and GFP signals in 50 HemSps were assessed.

**Extended Data Fig 3.**
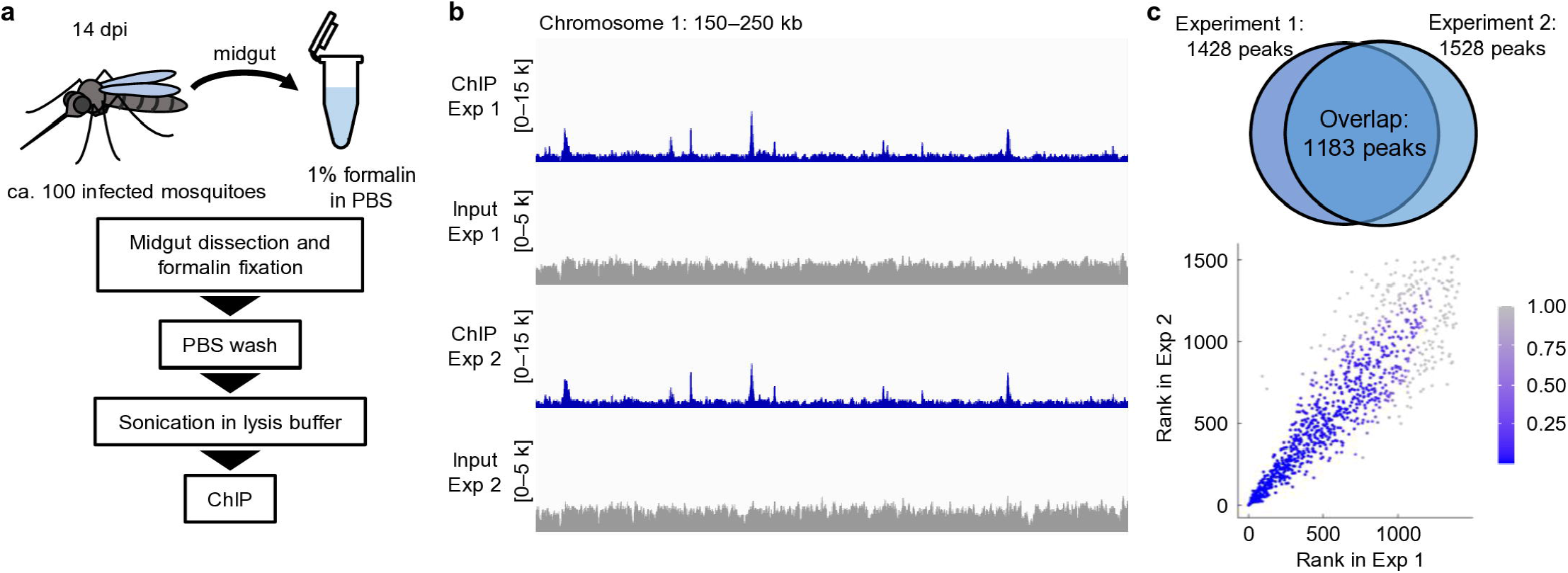
Chromatin immunoprecipitation followed by sequencing analysis of AP2-Sp3. (a) Schematic of chromatin immunoprecipitation (ChIP) experiment using AP2-Sp3::GFP at 14 dpi. (b) Integrative Genomics Viewer images for the duplicate AP2-Sp3 ChIP-seq experiments on a region of chromosome 1. ChIP (blue) and input (grey) data are shown. Histograms show the raw read coverage of ChIP data normalized by library size (bin size = 10 bp). Scales are indicated in square brackets. (c) Comparison of duplicate ChIP-seq experiments. Venn diagram shows the number of overlapping peaks between duplicates. Scatter plot shows IDR1D analysis comparing duplicates. The rank of peaks according to their *q* value for each experiment is plotted against each other. The Irreproducible Discovery Rate (IDR) of each peak was represented using a gradient color scale.

**Extended Data Fig 4.**
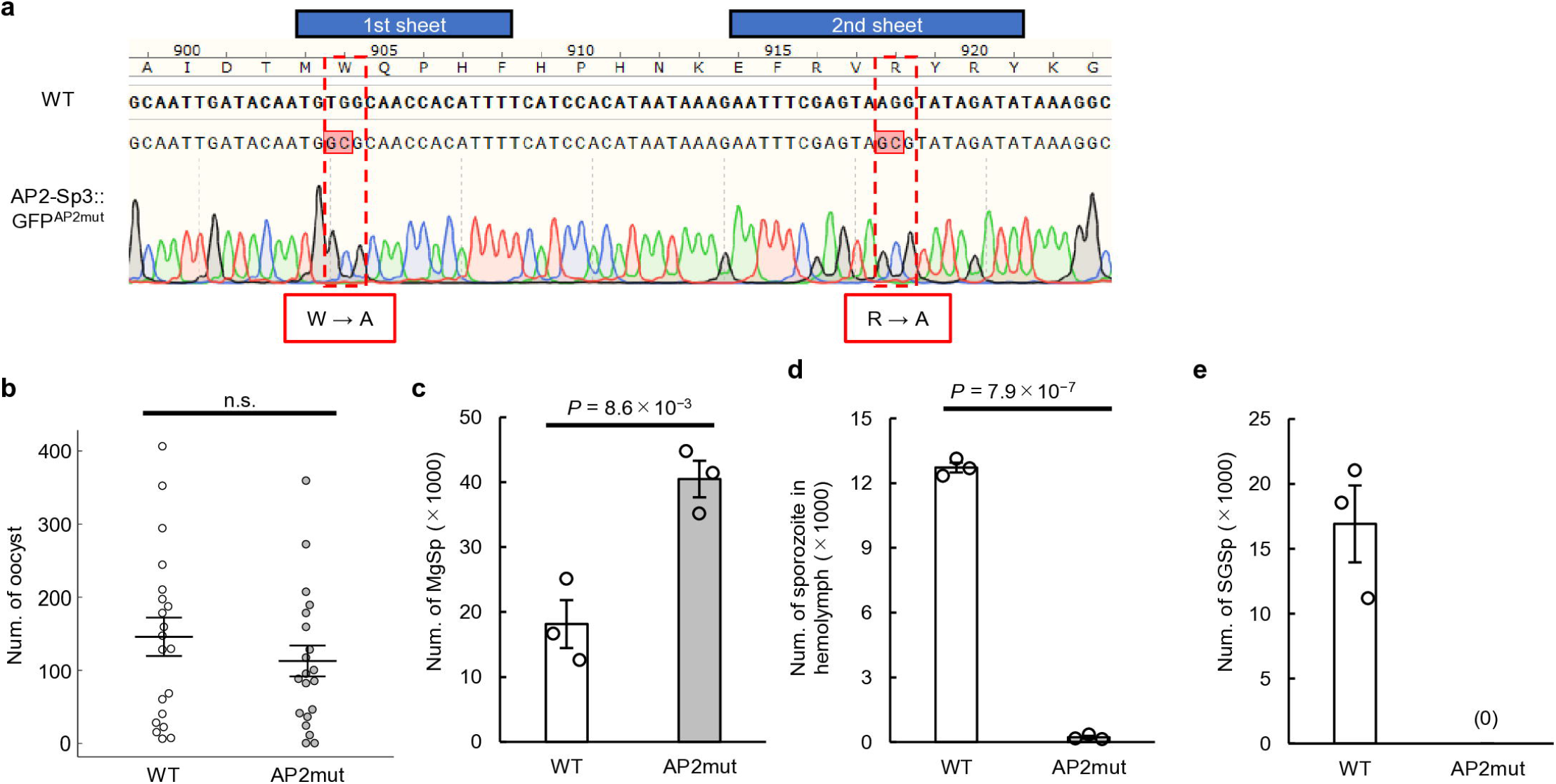
The AP2 domain of AP2-Sp3 is essential for its function. (a) Introduction of mutations into apicomplexan-conserved residues of the AP2-Sp3 AP2 domain (AP2-Sp3::GFP^AP2mut^). The Sanger sequence result after introducing mutations is shown under the wild-type sequence, and the resulting amino acid substitutions are indicated below. (b) The number of oocysts per mosquito infected with wild-type and AP2-Sp3::GFP^AP2mut^ at 14 dpi. Lines indicate the mean values and the standard error of the mean (SEM) (n = 20). n.s. indicates that the difference between mean values was not significant (a *P*-value calculated by two-tailed Student’s t-test was higher than 0.05.). (c) The number of midgut sporozoites per mosquito at 14 dpi. Ten midguts were pooled for each experiment. Error bars indicate the SEM from three biologically independent experiments. *P* values were calculated by two-tailed Student’s t-test. (d) The number of hemolymph sporozoites per mosquito at 20 dpi. Hemolymph from ten mosquitoes were pooled for each experiment. (e) The number of salivary gland sporozoites per mosquito at 21 dpi. Salivary glands from ten mosquitoes were pooled for each experiment. No SGSps were detected for AP2-Sp3::GFP^AP2mut^ [indicated as (0)].

**Extended Data Fig 5.**
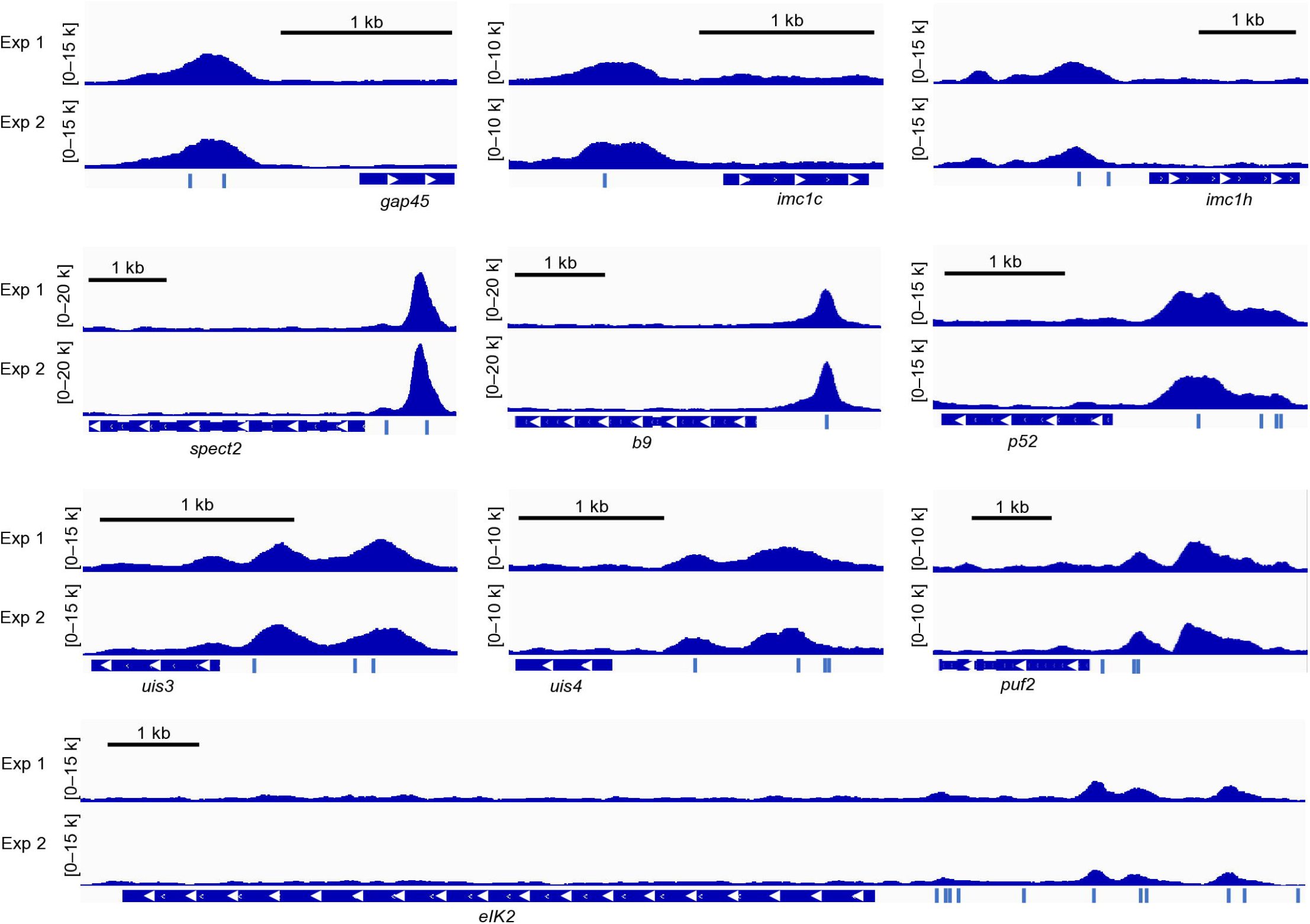
AP2-Sp3 targets include many sporozoite-related genes. Integrative Genomics Viewer showing AP2-Sp3 ChIP-seq peaks upstream its target genes (*gap45*, PBANKA_1437600; *imc1c*, PBANKA_1202000; *imc1h*, PBANKA_1436600; *spect2*, PBANKA_1006300; *b9*, PBANKA_0808100; *p52*, PBANKA_1002200; *uis3*, PBANKA_1400800; *uis4*, PBANKA_0501200; *puf2*, PBANKA_0719200; *eIK2*, PBANKA_0205800). Histograms show the raw read coverage of ChIP data normalized by library size (bin size = 10 bp). Scales are indicated in square brackets.

**Extended Data Fig 6.**
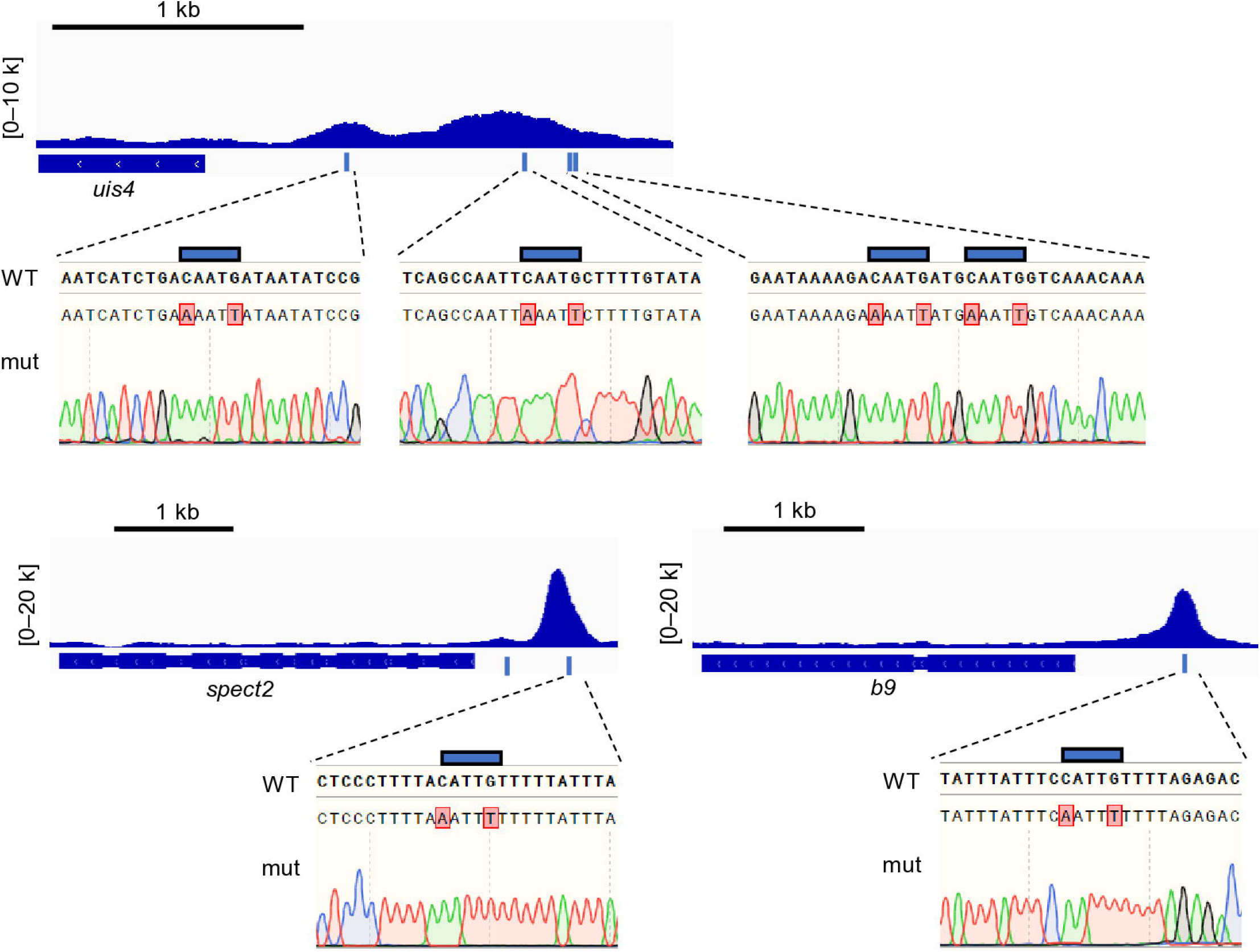
Introduction of mutations into AP2-Sp3-binding motif CATTG. IGV image showing AP2-Sp3 ChIP-seq peaks upstream of *uis4*, *b9*, and *spect2*. Genomic sequences around CATTG (indicated by a blue box) in the peak regions are shown with Sanger sequence results for mutant parasites, which carry mutations altering CATTG to aATTt.

**Extended Data Table 1.**
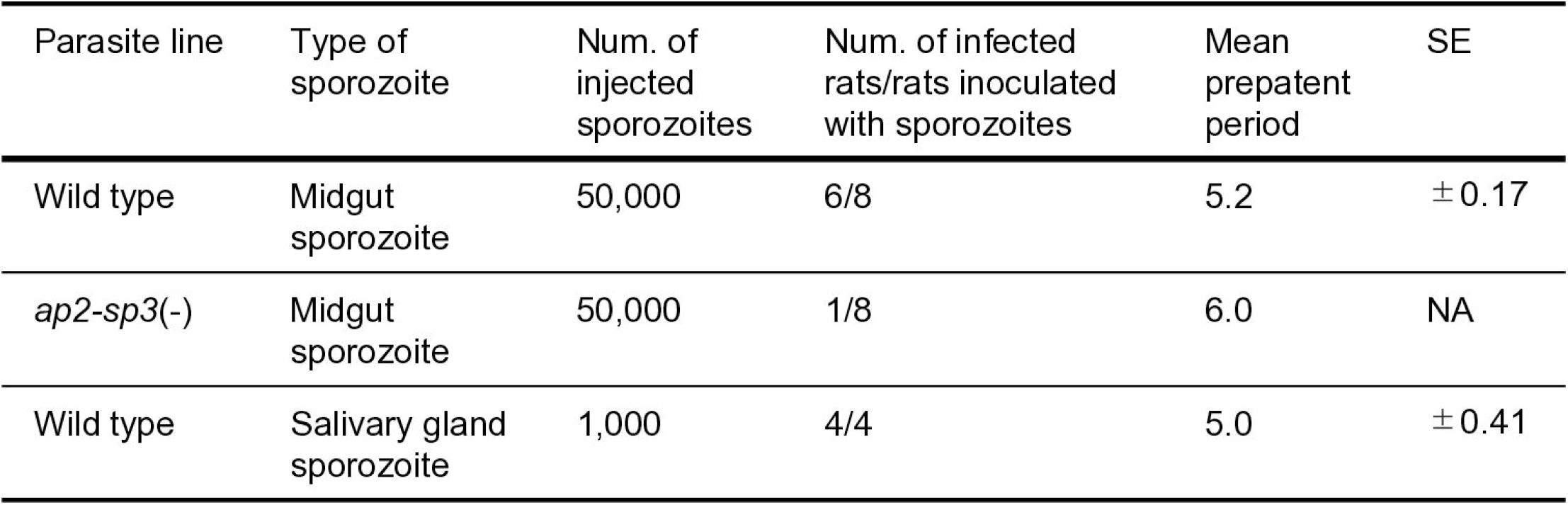
Infectivity assay of *ap2-sp3*(-) midgut sporozoites.

| Parasite line | Type of sporozoite | Num. of injected sporozoites | Num. of infected rats/rats inoculated with sporozoites | Mean prepatent period | SE |
| --- | --- | --- | --- | --- | --- |
| Wild type | Midgut sporozoite | 50,000 | 6/8 | 5.2 | $\pm 0.17$ |
| *ap2-sp3(-)* | Midgut sporozoite | 50,000 | 1/8 | 6.0 | NA |
| Wild type | Salivary gland sporozoite | 1,000 | 4/4 | 5.0 | $\pm 0.41$ |

