## Supplementary Figure for "Transcriptional repression by AP2-Sp3 regulates the mosquito-to-mammal infectivity switch in malaria sporozoites"

**Supplementary Information**

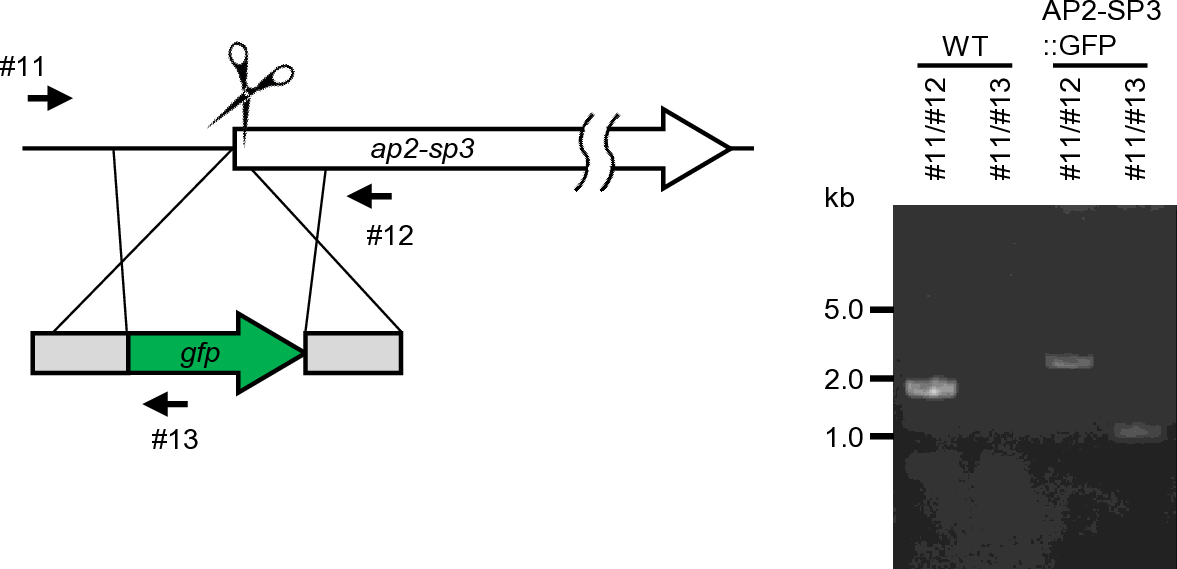

**Supplementary Fig 1. Genotyping of AP2-Sp3::GFP.**

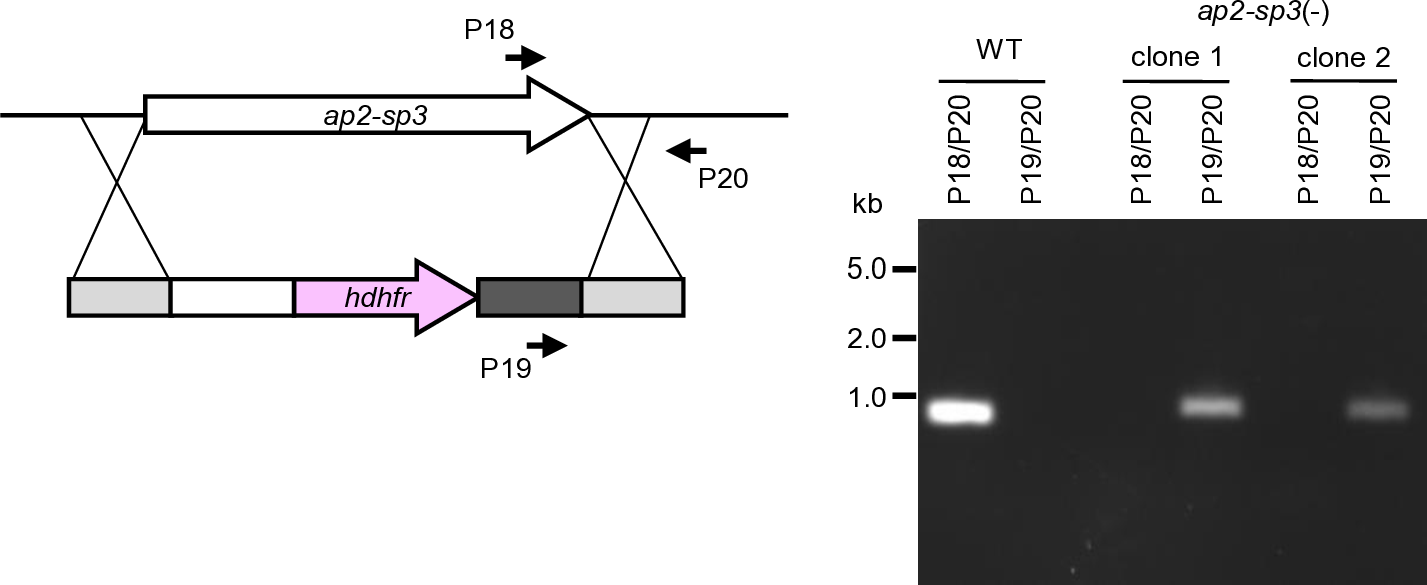

**Supplementary Fig 2. Genotyping of *ap2-sp3*(-).**

**Supplementary Table 1. Differential expression analysis between WT and *ap2-sp3*(-).**

**Supplementary Table 2. K-means clustering of 2000 most variable genes during sporozoite development.**

**Supplementary Table 3. ChIP-seq analysis of AP2-Sp3.**

(a) Peaks in experiment 1. (b) Peaks in experiment 2. (c) Motif enrichment analysis for 5-bp motifs. (d) Motif enrichment analysis for 6-bp motifs.

**Supplementary Table 4. ChIP-seq analysis of AP2-Sp3 with mutations within its AP2 domain.**

(a) ChIP-seq peaks. (b) Motif enrichment analysis for 5-bp motifs. (c) Motif enrichment analysis for 6-bp motifs.

**Supplementary Data 5. Target genes of AP2-Sp3.**

(a) List of targets. (b) Gene ontology analysis.

**Supplementary Table 6. List of primers used in this study.**
